# FQSqueezer: *k*-mer-based compression of sequencing data

**DOI:** 10.1101/559807

**Authors:** Sebastian Deorowicz

## Abstract

**Motivation:** The amount of genomic data that needs to be stored is huge. Therefore it is not surprising that a lot of work has been done in the field of specialized data compression of FASTQ files. The existing algorithms are, however, still imperfect and the best tools produce quite large archives.

**Results:** We present FQSqueezer, a novel compression algorithm for sequencing data able to process single- and paired-end reads of variable lengths. It is based on the ideas from the famous prediction by partial matching and dynamic Markov coder algorithms known from the general-purpose-compressors world. The compression ratios are often tens of percent better than offered by the state-of-the-art tools.

**Availability and Implementation:** https://github.com/refresh-bio/fqsqueezer

**Supplementary information:** Supplementary data are available at publisher’s Web site.

## 1 Introduction

In the recent years, genome sequencing has became a mature technology with numerous applications in the medicine. The instruments by Illumina (encountered to the 2nd generation of sequencers) produce majority of available data for little money (e.g., about one thousand of U.S. dollars for human whole genome sequencing). The reads are relatively short (up to a few hundreds of bases) but are of high quality. The 3rd generation instruments by PacBio or Oxford Nanopore can deliver much longer reads but unfortunately of much worse quality or at much lower throughput.

In studies like (Stephens *et al.*, 2015; Deorowicz & Grabowski, 2013), the estimations of the amount of genomic data and the related costs can be found. The presented huge numbers directly lead to the conclusion that in the near future just a storage and transfer of sequenced (and mapped) reads will consume a lot of money and could be a dominant factor in total costs related to sequencing. Therefore, it is not surprising that a lot of research was made in this field. The obvious first step was application of some general-purpose compressor, like gzip. The about 3-fold reduction of files was remarkable, but the data deluge asked for more. The main problem of gzip is that it was designed mainly for textual data, or more precisely, for data with similar redundancy types. Unfortunately things like repetitions of parts of data at short distances are uncommon in read collections.

The next step was an invention of specialized algorithms taking into account types of redundancies specific for FASTQ files (Cock *et al.*, 2010). Some of the most important early results were presented in (Deorowicz & Grabowski, 2011; Hach *et al.*, 2012; Bonfield and Mahoney, 2013; Roguski and Deorowicz, 2014). The details of the proposed algorithms were different, but in general the authors tried (with some exceptions) to compress the reads locally, i.e., not looking for the large-scale relations between the reads. The reason was rather simple and practical: the amounts of memory necessary to construct a dictionary data structure allowing to find overlaps between reads could be a few times larger than the input file size, e.g., hundreds of GB for human genomes. The improvement over gzip was, however, limited. For example, the best algorithms were able to reduce the space necessary for DNA bases about 5 times. This value could be compared to 4-fold reduction by simple spending 2 bits to distinguish between all 4 valid bases. Moreover, it appeared that the compression of quality values was even more problematic.

This situation motivated researchers to look for alternatives. At the beginning, they focused just on the compression of bases. The key idea was to reorder the data to gather reads originating from close regions of genomes. This could seem as a loss of information, but as the original ordering of reads in a FASTQ file is usually more or less random and one can argue that it is hard to say which of two random orderings is better (and even how we can define what “better” means here). The first notable attempt into this direction was the work by Cox *et al.* (2012). The authors introduced of a variant of the Burrows–Wheeler transform to find overlaps between reads. For human reads with 40-fold coverage they were able to spend about 0.5 bits per base, which was a significant improvement.

In the following years, other researchers explored the concept of using minimizers (Roberts *et al.*, 2004), i.e., short, lexicographically smallest, substrings of sequences, to find reads from close regions. The key observation was that if two reads originate from the close regions of a genome their minimizers are usually the same. Thus, the reads can be grouped by their minimizers. In the first work following this idea (Grabowski *et al.*, 2015), for the mentioned human dataset it was sufficient to use just a bit more than 0.3 bits per base. A similar result was obtained later in (Patro and Kingsford, 2015). The possible gains were, however, limited by the fact that the reads identified to originate from the same genome region could span no more than two read lengths.

In (Roguski *et al.*, 2018), it was shown how to group reads from a bit larger genome regions. Significantly better results were, however, obtained in three recent articles presenting HARC (Chandak *et al.*, 2018a), Spring (Chandak *et al.*, 2018b), and Minicom (Liu *et al.*, 2018). The attempts differ in details, but are based on similar ideas. The overlaps are found for much larger genome regions (in theory up to chromosome size) thanks to dictionary structures storing minimizers of parts of reads. What is also worth to mention, FaStore and Spring do not focus just on DNA bases and they can compress also the complete FASTQ files.

Together with improving the compression ratio for DNA symbols, the quality scores became responsible for a dominant part of the compressed archives. Therefore a number of works focused on this problem. One of the simplest strategies was to reduce the resolution of quality scores. Illumina in their HiSeq sequencers restricted the quality scores to eight values, and then in the NovaSeq instruments to just four values. The rational for these decisions was that the quality of sequencing is currently very good and the prices of sequencing are low. Therefore, if necessary, it is easier (and cheaper) to perform sequencing with a bit larger coverage than store high-resolution quality scores. The recent experiments suggest that reduction of quality score resolution has very little (if any) impact on the quality of variant calling. For example, in (Roguski *et al.*, 2018) it was shown that even more aggressive reduction to just two quality values can be justified at least in some situations. Moreover, there are several algorithms like QVZ (Malysa *et al.*, 2015; Hernaez *et al.*, 2016), Crumble (Bonfield *et al.*, 2018) that perform advanced analysis of quality scores to preserve only the most important information.

In this article, we propose a novel compression algorithm for FASTQ files. The main novelty is in the compression of DNA bases, as for quality scores and read identifiers we follow similar strategies as in the top existing tools. Our algorithm, FQSqueezer, is based on the ideas from the prediction by partial matching (PPM) (Cleary and Witten, 1984; Moffat, 1990) and dynamic Markov coder (DMC) (Cormack & Horspool, 1987) general-purpose methods. A direct adaptation of the PPM-like strategy to sequencing reads would be, however, very hard and likely unsuccessful. There are at least four main reasons for that. First, in the ideal case, the PPM algorithm should construct a dictionary of all already seen strings of length up to some threshold, significantly larger than log_4_(*genome*_*size*), which for human genomes seems to be unimplementable on workstations and even medium-sized servers. Second, the PPM algorithms often need many accesses to the main memory to compress a single symbol. For huge dictionaries this could result in a very slow processing (due to cache-misses). Third, sequencing data contain errors that should be corrected to refrain from expansion of the dictionary structures. Fourth, the PPM algorithms usually learn slowly, which is a good strategy for texts, but seems to be bad for genomic data.

To overcome these problems we designed a few fixed-*k* dictionaries for *k*-mers found in the reads. Moreover, the dictionaries are organized in a way reducing the number of cache misses. We also estimate the probability of symbols occurrence much more aggressively, which results in significantly better compression (compared to classical PPM-like estimation). Finally, just for the storage of *k*-mers in the dictionaries we perform some kind of error correction.

The main asset of FQSqueezer is its compression ratio, usually much better than of the state-of-the-art competitors, i.e., FaStore (Roguski *et al.*, 2018), Spring (Chandak *et al.*, 2018b), and Minicom (Liu *et al.*, 2018). Our tool has, however, also some drawbacks in terms of speed and memory usage. Namely, it is a few times slower than the mentioned competitors in compression and much slower in decompression.

## 2 Methods

### 2.1 Basic definitions

It will be convenient to start from a definition of some terms. Let *x* = *x*_1_*x*_2_ *… x*_*r*_ be a sequence of symbols from alphabet {A, C, G, T, N}. The *length* (*size*) of a sequence will be the number of elements it is composed of. A *substring* can be obtained from a sequence by removing (possibly 0) symbols from the beginning and the end. The notation *x*_*i,j*_ means a sub-string *x*_*i*_*x*_*i*+1_ *… x*_*j*_. A *k-mer* is a sequence of length *k*. A *canonical k*-mer is lexicographically smaller of a *k*-mer and its reverse complement. A *kernel* of a *k*-mer is a *k*-mer subtracted by two first and two last symbols.

### 2.2 Basic description of the algorithm

FQSqueezer is a multi-threaded algorithm, but for simplicity of presentation we will start from a single-threaded variant. Out tool accepts both single-end (SE) and paired-end (PE) reads. The reads can be stored in the original ordering (OO) or can be reordered (REO). In the reordering mode, the reads are initially sorted according to the DNA sequence (first read of a pair in the PE mode).

The input FASTQ files (or sorted files in the REO mode) are loaded in blocks of size 16MB. The reads from a single block (pair of blocks in the PE mode) are compressed one by one (or pair by pair). The read ID and quality values are compressed using rather standard means (similarly like in the top existing FASTQ compressors). The details are described in Sections 2.5 and 2.6. Below, we will focus just on the DNA symbols.

The processing of a SE read (or first read of a pair) is composed of compressing of a prefix and a suffix. For each base we determine the statistics of occurrences of *k*-mers ending at this base in the already processed part of the input data. To this end, we maintain a few dictionaries: *D*_e_, *D*_p_, *D*_s_, and *D*_b_ that store numbers of occurrences of: *e*-mers, *p*-mers, *s*-mers, and *b*-mers, respectively, where *e < p < s < b*. The details of the organization of the dictionaries are given in Section 2.3.

In general, the longest possible, but no longer than *b* − 1 symbols, context (substring preceding the current symbol in a read) is taken to predict the current symbol. Then, the symbol is encoded using these predictions (as well as some other properties of a read and the current position) with a use of range coder. The details are given in Section 2.4.

In the REO mode, the prefix of size *p* of a SE read (or first read of a pair in the PE mode) is encoded differently. Roughly speaking, we treat it as a 2*p*-bit unsigned integer and encode the difference between it and the integer representing the prefix of the previous read. The suffix is compressed in the same way as in the OO mode.

The second read of a pair is compressed in a bit more complex manner, but the processing is the same in the OO and REO modes. Initially we try to predict some its *b*-mer. To this end we use a dictionary *M*_b_ that stores pairs of minimizers of read pairs seen so far. Quite often this allows to encode the *b*-mer of the second read in an efficient way. Then, we store the substrings of the read following and preceding this *b*-mer. If we were unable to predict the minimizer of the second read, we store the read in the same way as a SE read in the OO mode.

### 2.3 Data structures organization

The dictionaries are organized in various ways. The *D*_p_ dictionary is a vector of 2-bit integers of size 4^*p*^. It is indexed by an integer representation of a *p*-mer. Each 2-bit field stores a number of occurrences of a related *p*-mer, saturated at 3. When a *p*-mer is added to the dictionary the counters are incremented for both *p*-mer and its reversed complement.

The *D*_s_ dictionary is a collection of 4^*s*−10^ hash tables representing canonical *s*-mers and the counters (no. of occurrences of each *s*-mer). When looking for (or inserting to) *s*-mer *x* in the dictionary, a hash table from the collection is identified by the substring *x*_3,*s*−8_. Each 32-bit entry of a hash table contains: 4 bits for *x*_1,2_, 16 bits for *x*_*s*−7,*s*_, and 12 bits for the counter. A hash function calculates its value just using *x*_3,*s*−2_, i.e., the kernel of an *s*-mer. Such an organization allows to make use of software prefetching mechanism to reduce delays related to cache misses. Moreover, the *s*-mers sharing their (*s* − 1) starting symbols are at close positions, which also reduces the number of cache misses. The maximal value of a counter is 2^12^ −1, which can be saturated for repetitive genomes. Therefore, the counter incrementation is a bit tricky. Counter values *v* not exceeding 2^11^ are always incremented. The larger values are incremented with probability 1*/*(*v* − 2^11^). Thus, the large counts are approximate, but the maximal representable value is much larger.

The *D*_b_ dictionary is organized in a similar way as *D*_s_, but here we have a collection of 4^*b*−13^ hash tables. The hash table is identified by an integer representation of *x*_3,*b*−11_. The 32-bit entries are composed of: 4 bits for *x*_1,2_, 22 bits for *x*_*b*−10,*b*_, and 6 bits for the counter. The incrementation of counter value *v* larger than 8 is with probability 1*/*2(*v* − 8).

The compressor maintains global dictionaries *D*_p_, *D*_s_, and *D*_b_ that can be queried in parallel by separate threads. The update of these dictionaries is after processing the complete 16 MB blocks (in fact for the 1st, 2nd,…, 99th block it is more frequent, i.e., 100 *- block*_*number* times for a block). It is made by all threads as each thread updates different hash tables from a collection (or different part of vector *D*_p_). The delayed update means, however, that the recently processed *k*-mers are absent from the global dictionaries. To overcome this problem, each thread stores also small local dictionaries 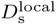 and 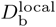 for representation of *s*-mers and *b*-mers seen in this thread from the last update of the global dictionaries. This local dictionaries are hash tables with 64-bit entries.

The *D*_e_ dictionary is a hash table storing statistics of occurrences of all substrings of length from 1 to *e*.

In the PE mode, we need also *M*_b_ dictionary. It is a hash table containing 128-bit entries storing two *b*-mers that are signatures (variant of a minimizer, defined as in (Deorowicz *et al.*, 2015)) of some parts of reads and the counter stored at 64 - 2*b* bits. More precisely, for each pair of reads *x* and *y* we determine several signatures, i.e.,

- <u>*x*^1^</u>, <u>*x*^2^</u>, <u>*x*^3^</u> — the signatures in the first, second, and third part of *x*,
- <u>*y*^1^</u>, <u>*y*^2^</u>, <u>*y*^3^</u> — as above for *y*,
- <u>*x*^23rc^</u> and <u>*y*^23rc^</u> — the signatures in the reverse-complemented second and third part of *x*, and *y*.

Then, we add the following pairs to *M*_b_: ⟨*<u>x</u>*<u>^1^</u>, *<u>y</u>*<u>^1^</u>⟩, ⟨*<u>x</u>*<u>^1^</u>, *<u>y</u>*<u>^3^</u>⟩, ⟨*<u>x</u>*<u>^1^</u>, *<u>x</u>*<u>^23rc^</u>⟩, ⟨*<u>x</u>*<u>^2^</u>, *<u>y</u>*<u>^1^</u>⟩, ⟨*<u>x</u>*<u>^2^</u>, *<u>y</u>*<u>^3^</u>⟩, ⟨*<u>x</u>*<u>^3^</u>, *<u>y</u>*<u>^1^</u>⟩, ⟨*<u>x</u>*<u>^3^</u>, *<u>y</u>*<u>^3^</u>⟩, ⟨*<u>y</u>*<u>^1^</u>, *<u>x</u>*<u>^1^</u>⟩, ⟨*<u>y</u>*<u>^1^</u>, *<u>x</u>*<u>^3^</u>⟩, ⟨*<u>y</u>*<u>^1^</u>, *<u>y</u>*<u>^23rc^</u>⟩, ⟨*<u>y</u>*<u>^2^</u>, *<u>x</u>*<u>^1^</u>⟩, ⟨*<u>y</u>*<u>^2^</u>, *<u>x</u>*<u>^3^</u>⟩, ⟨*<u>y</u>*<u>^3^</u>, *<u>x</u>*<u>^1^</u>⟩, ⟨*<u>y</u>*<u>^3^</u>, *<u>x</u>*<u>^3^</u>⟩. Th local dictionary 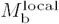 is maintained in the similar way as local variants of other dictionaries.

The choice of parameters *f*, *p, s, b* has some impact on the compression ratios. They should be adjusted to the genome size of the sequenced organism (or a sum of genome sizes in metagenomic experiments). Therefore, the user should specify the genome size, which is used to set the parameters as shown in Table 1. If the genome size is unknown, it roughly can be guessed from the FASTQ file size and the sequenced coverage.

**Table 1.**
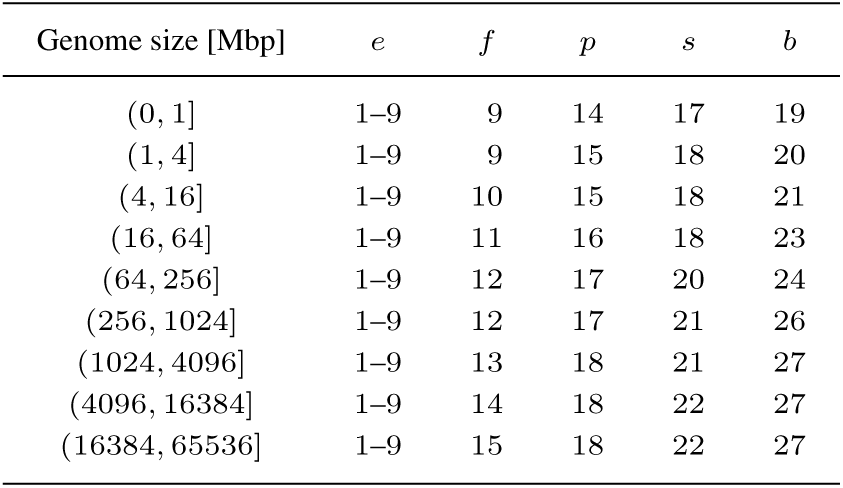
Determination of dictionary parameters

| Genome size [Mbp] | $e$ | $f$ | $p$ | $s$ | $b$ |
| --- | --- | --- | --- | --- | --- |
| (0, 1] | 1–9 | 9 | 14 | 17 | 19 |
| (1, 4] | 1–9 | 9 | 15 | 18 | 20 |
| (4, 16] | 1–9 | 10 | 15 | 18 | 21 |
| (16, 64] | 1–9 | 11 | 16 | 18 | 23 |
| (64, 256] | 1–9 | 12 | 17 | 20 | 24 |
| (256, 1024] | 1–9 | 12 | 17 | 21 | 26 |
| (1024, 4096] | 1–9 | 13 | 18 | 21 | 27 |
| (4096, 16384] | 1–9 | 14 | 18 | 22 | 27 |
| (16384, 65536] | 1–9 | 15 | 18 | 22 | 27 |

### 2.4 Algorithmic details

There are 4 different modes in which FQSqueezer can be run: (*i*) for single-end (SE) reads with preserving the original ordering of the reads, (*ii*) for SE reads without preserving the original ordering of the reads, (*iii*) for paired-end (PE) reads with preserving the original ordering of the reads, (*iv*) for PE reads without preserving the original ordering of the reads. Most of the algorithmic details are the same in the modes, so we will start the description from the simplest SE order-preserving mode.

#### SE reads — original ordering

The first *f* symbols of a read are compressed using the statistics of occurrence of up to *e* previous symbols. This value is 1 at the beginning of compression and quite fast grows to 9 (when there are sufficient amount of statistics). The processing of successive symbols is as follows. We maintain 3 pairs of *k*-mers:

- (*p* − 1)-mers: 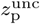 and *z*_p_, where the first contains recently seen *p*−1 symbols and the second is a copy of 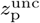 in which some symbols could be changed by our correction mechanism,
- (*s* − 1)-mers: 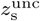 and *z*_s_ defined as above for *s* − 1 recent symbols,
- (*b* − 1)-mers: 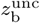 and *z*_b_ defined as above for *b* − 1 recent symbols.

At the beginning of the processing of a read the *k*-mers are incomplete.

Let us assume that we are at the *i*th position in a read *x*. First, we try to obtain as accurate predictions of the current symbol as possible. To this end, if *s < i* we ask *D*_b_ and 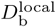 for statistics of occurrences of *z*_b_ followed by A, C, G, or T. If *z*_b_ is complete, the queries are straightforward. Otherwise, to collect the statistics we fill *z*_b_ (at the beginning) by all possible combinations of bases, which can be costly (in terms of time). In case of no success, we try also the same for 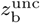.

If we were unable to collect any statistics and *p < i*, we try the similar procedure for *z*_s_ and *D*_s_, 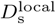. Moreover, we use this step also in situation in which for more than one base the counters obtained from querying *D*_b_ achieve 63, the maximal counter value.

If necessary (no success), the search is performed for *z*_p_ and *D*_p_.

If we were not able to find any statistics in the previous stages we perform more aggressive search. If *b ≤ i*, we try to collect the statistics for *z*_b_ and *D*_b_ for approximate matches (with 1 mismatch). Unfortunately, this means dozens of queries. In case of no success and *s ≤ i*, we try the same for *z*_s_ and *D*_s_. Finally, if necessary, we use also *z*_p_ and *D*_p_.

If after this search procedure we do not have any statistics, we encode the current symbol taking into account from 1 to 9 previous symbols (exactly like for the read prefix). The situation is more interesting if we were able to collect some statistics. We form a context for encoding of the current symbol. The context components are:

1. *pos* — current position in a read,
2. *pattern* — sorted counters of occurrences of symbols,
3. *cnt_lev* — origin of statistics, one of:

- *z*_b_ or 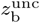 and *D*_b_ or 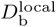,
- *z*_s_ or 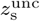 and *D*_s_ or 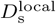,
- *z*_p_ or 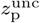 and *D*_p_,
- *z*_b_ or 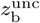 and *D*_b_ or 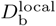 with additional information from *z*_*s*_ and *D*_*s*_,
4. *accuracy* — number of accurate predictions of the actual at previous 8 positions,
5. *top_symbol* — the symbol with the highest counter value,
6. *correction* — one of:

- there was a correction in the most recent half of *z*_b_,
- there was a correction in the former half of *z*_b_,
- the statistics are from the approximate queries,
- none of the above.

The actual context is constructed from the context components in the way motivated by the dynamic Markov coder (Cormack & Horspool, 1987). We allow 7 levels of contexts. In the parentheses we give the number of times the context can be used before it is split into more detailed ones (i.e., before we go to the next level):

1. context is empty (1),
2. context contains (32):

- *cnt_lev*,
- *pos* rounded to 16 if we are not at the beginning of a read (values smaller than *b* are not rounded) and not close to the end of a read (last 5 positions in a read are stored explicitly),
- two largest counters down-sampled to: 30 values (if the counters are from the *D*_p_ queries) or 10 values (otherwise),
- the remaining counters down-sampled to: 10 values (if the counters are from the *D*_p_ queries) or 5 values (otherwise),
3. context contains additionally: *accuracy* and *correction* (64),
4. if the counters are not obtained from the *D*_p_, the two largest counters are down-sampled to 15 values (64),
5. if the counters are not obtained from the *D*_p_, the two largest counters are down-sampled to 30 values and the remaining counters to 10 values, otherwise the two smallest counters are down-sampled to 30 values (128),
6. context contains additionally: *top_symbol* (512),
7. the positions rounded to 16 are now rounded to 8.

At the end of processing of a current position, we try to correct (just to make better representation in the dictionaries) the current symbol. If *i < b* we do nothing with the exception for a case in which the current symbol is N, as we update it to A. For *i ≥ b* we alternatively run two correction procedures. First is applied if the counters are from inspecting the *D*_b_ or 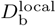. If the counter value for the current symbol is 0 and the most frequent symbol has counter 3 or more we make a substitution. Second is applied in case of no statistics or if the statistics are from the *D*_p_. Moreover, to apply this correction the current coverage (estimated from the number of updates to *D*_p_) must be at least 7. We try to change one of the previous 6 symbols to find a match in the *D*_b_.

As a simple time optimization at the beginning of processing of a read we check whether it is a copy of the previous read. We store this information and do not process copied reads position by position.

#### SE reads — reordered

In the REO mode, the reads are initially split into 256 bins according to the first 4 symbols (Ns are treated as Ts). Then the reads are read from bins and sorted in the main memory before further processing.

The only difference in the compression from the OO mode is handling of the read prefix. We treat the first *p* symbols as an 2*p*-bit unsigned integer *t*. Let *t*^prev^ be such value for the previous read. We check in *D*_p_ the counter *c* for *t*. We encode *c* and then determine (in *D*_p_) the number of *p*-mers between *t*^prev^ and *t* with counter equal *c* and store this number.

A special care is necessary for Ns in the read prefix. They are replaced by Ts for the conversion to the integers, but a special flag is stored to tell whether such substitutions took place in the present read. If necessary the substitutions are stored to allow perfect reconstruction of the prefix.

#### PE reads — original ordering

The compression of PE reads in the OO mode is similar to the compression of SE reads. The first read of a pair is compressed exactly in the same way. The second read also can be compressed in the same way, but fortunately, quite often we can do much better. To this end, we determine 4 signatures of the first read, i.e., form the 1st, 2nd, 3rd, 4th part of the read. For each signature we ask *M*_b_ for the paired candidate signatures. We merge, the lists and pick the top 16 candidates. Then in the second read of a pair we look for the candidates. In case of success, we just store the position of the found signature at the candidate list and the position of the signature in the second read. Then, we store the parts after and before the signature in the second read using the same way as in the SE mode.

#### PE reads — reordered

The compression of PE reads in the REO mode is a mix of the compression of SE-REO and PE-OO modes. The first read of a pair is stored as in the SE-REO mode. The second one is processed as in the PE-OO mode.

### 2.5 Compression of IDs

In the lossless mode, IDs are compressed similarly like in the top compressors, like Spring or FaStore. The ID of each read is tokenized (the separators are non-alphanumerical characters). Then the tokens are compared with the tokens of the previous read. If the tokens at corresponding positions contain numerical values, the difference between the integers is calculated and stored. Otherwise the corresponding tokens are compared as strings. If they are equal we just store a flag. In the opposite case, we store a mismatch flag and compare the tokens character by character storing the result of comparison and (if necessary) the letter from the current read. It can also happen that the list of tokens differ significantly, i.e., they are of different length or the corresponding tokens are of different type. In this situation the ID is stored character by character.

FQSqueezer offers also two lossy modes. In the first one, it preserves just the instrument name (first part of the ID). These names are organized in a move-to-front list (McCabe, 1965) and the position of the current ID at the list is encoded. If the current instrument name is absent form the list, it is encoded explicitly. In the last possible mode, IDs are discarded.

### 2.6 Compression of quality scores

The quality scores can be compressed in five modes allowing different number of values: 64, 8, 4, 2, none. If the input FASTQ file has already reduced resolution of quality scores (e.g., 4 for Illumina NovaSeq sequencers), no conversion is necessary. Otherwise, in the lossy mode the necessary resolution reduction is made by the compressor. The encoding is made using contexts containing the position in a read and 2 (64-value alphabet), 6 (8-value alphabet), 9 (4-value alphabet), or 10 (binary alphabet) previous scores.

### 2.7 Implementation details

The implementation is in the C++14 programming language. All hash tables use linear probing for collision handling. To reduce delays of cache misses we make use of software prefetching, e.g., before processing the current symbol we know the kernels of maintained *k*-mers, so we can prefetch dictionaries. The multithreading is implemented using the native C++ threads. The FASTQ blocks are split into as many parts as the number of threads. Each thread processes its part independently and the global dictionaries are available only for querying. At the synchronization points the threads update the global dictionaries. Nevertheless, the threads update different parts of the dictionaries so locks are unnecessary.

## 3 Results

### 3.1 Tools and datasets

For the evaluation we used the state-of-the-art competitors, i.e., FaStore (Roguski *et al.*, 2018), Spring (Chandak *et al.*, 2018b), and Minicom (Liu *et al.*, 2018). We resigned from testing some other good compressors like BEETL (Cox *et al.*, 2012), Orcom (Grabowski *et al.*, 2015), HARC (Chandak *et al.*, 2018a), AsembleTrie (Ginart *et al.*, 2018) as the previous works demonstrated that they perform worse than the picked tools. The older utilities are not competitive in terms of compression ratio. Some of them are very fast, e.g., DSRC 2 (Roguski and Deorowicz, 2014). Nevertheless, in this article we focus mainly on the compression ratio.

The datasets for experiments are taken from the previous studies. They are characterized in Supplementary Table 1. In the main part of the article, we used 9 datasets but more results can be found in the Supplementary Worksheet.

All experiments were run at workstation equipped with two Intel Xeon E5-2670 v3 CPUs (2 *×* 12 double-threaded 2.3GHz cores), 256GB of RAM, and six 1 TB HDDs in RAID-5. If not stated explicitly the programs were run with 12 threads.

### 3.2 Compression of the bases

The most important part of the present work is the compression of bases. Therefore, in the first experiment we evaluated the tools in this scenario. The results for the SE reads are given in Table 2. The ratios are in output bits per input base. As it is easy to observe, in the majority of cases FQSqueezer outperforms the competitors. For some datasets the gain is large. Nevertheless, for two datasets FQSqueezer looses to Minicom in the OO mode. Probably the case is that the reads in this dataset are not at random ordering and Minicom can benefit from this. The results for the REO mode suggest to confirm this supposition.

**Table 2.** Compression ratios for single-end reads

| Dataset | Size<br>[Gbp] | Original ordering |  |  |  | Reordered |  |  |  |  |
| --- | --- | --- | --- | --- | --- | --- | --- | --- | --- | --- |
|  |  | Spring | Minicom | FQSqueezer | Gain | FaStore | Spring | Minicom | FQSqueezer | Gain |
| ERR174310_1 | 20.97 | 0.696 | 0.857 | **0.649** | 7.4 | 0.593 | 0.408 | 0.589 | **0.396** | 3.1 |
| ERR532393_1 | 3.58 | 0.667 | 0.647 | **0.528** | 22.5 | 0.482 | 0.433 | 0.410 | **0.294** | 39.3 |
| SRR327342_1 | 0.95 | 0.524 | 0.538 | **0.488** | 7.4 | 0.267 | 0.158 | 0.166 | **0.142** | 11.3 |
| SRR554369_1 | 0.17 | 0.441 | 0.509 | **0.414** | 6.6 | 0.494 | 0.240 | 0.307 | **0.227** | 5.6 |
| SRR635193_1 | 1.47 | 0.697 | 0.702 | **0.633** | 10.0 | 0.333 | 0.267 | 0.289 | **0.216** | 23.9 |
| SRR689233_1 | 1.48 | 0.449 | 0.442 | **0.400** | 10.6 | 0.248 | 0.193 | 0.187 | **0.149** | 25.5 |
| SRR870667_1 | 7.48 | 1.460 | 1.364 | **0.721** | 89.3 | 0.722 | 1.292 | 1.212 | **0.506** | 42.6 |
| SRR1265495_1 | 1.70 | 0.632 | **0.448** | 0.506 | −11.4 | 0.368 | 0.500 | 0.319 | **0.246** | 29.7 |
| SRR1265496_1 | 1.48 | 0.646 | **0.484** | 0.517 | −6.4 | 0.391 | 0.507 | 0.352 | **0.264** | 33.4 |

Compression ratios are in output bits per base [bpb]. Best results are in bold. ‘Gain’ (expressed in %) is defined as: best competitor ratio divided by FQSqueezer ratio subtracted by 1. FaStore does not offer original ordering preserving.

The results for the PE reads can be found in Table 3. In the OO mode FQSqueezer usually outperforms Spring significantly. Nevertheless, for the largest dataset it looses slightly. It is hard to say what is the reason. The situation for the REO mode is similar, but the gains are usually even larger than in the OO mode.

**Table 3.** Compression ratios for paired-end reads

| Dataset | Size<br>[Gbp] | Original ordering |  |  | Reordered |  |  |  |  |
| --- | --- | --- | --- | --- | --- | --- | --- | --- | --- |
|  |  | Spring | FQSqueezer | Gain | FaStore | Spring | Minicom | FQSqueezer | Gain |
| ERR174310 | 41.93 | **0.463** | 0.474 | −2.3 | 0.844 | **0.317** | 0.481 | 0.346 | −8.3 |
| ERR532393 | 7.15 | 0.623 | **0.462** | 34.7 | 0.694 | 0.505 | 0.471 | **0.347** | 35.8 |
| SRR327342 | 2.08 | 0.490 | **0.355** | 38.0 | — | 0.307 | — | **0.197** | 55.7 |
| SRR554369 | 0.33 | 0.322 | **0.299** | 7.7 | 0.652 | 0.222 | 0.293 | **0.205** | 8.4 |
| SRR635193 | 2.94 | 0.562 | **0.501** | 12.2 | 0.500 | 0.339 | 0.419 | **0.292** | 16.0 |
| SRR689233 | 2.96 | 0.367 | **0.309** | 18.8 | 0.350 | 0.239 | 0.257 | **0.184** | 30.2 |
| SRR870667 | 12.60 | 1.070 | **0.553** | 93.3 | — | 0.928 | — | **0.424** | 119.0 |
| SRR1265495 | 3.40 | 0.484 | **0.373** | 29.7 | 0.437 | 0.436 | 0.316 | **0.228** | 38.2 |
| SRR1265496 | 2.96 | 0.500 | **0.389** | 28.4 | 0.479 | 0.449 | 0.344 | **0.247** | 39.0 |

Compression ratios are in output bits per base [bpb]. Best results are in bold. ‘Gain’ (expressed in %) is defined as: best competitor ratio divided by FQSqueezer ratio subtracted by 1. FaStore and Minicom do not support original ordering preserving. For two datasets (REO mode) we were unable to run FaStore and Minicom.

The most important drawbacks of FQSqueezer are, however, its time and space requirements (Table 4 and Supplementary Worksheet). In the compression, it is a few times slower than the competitors, but in the decompression the difference is larger. The reason is simple. FaStore, Spring, Minicom need time to find overlapping reads that likely origin from close genome positions. Nevertheless, decoding of the matches between the overlapping reads is very fast. FQSqueezer can be classified as a PPM algorithm. In the decompression, the algorithms from this family essentially mimic the same work made in the compression, so the differences in compression and decompression times are negligible.

**Table 4.**
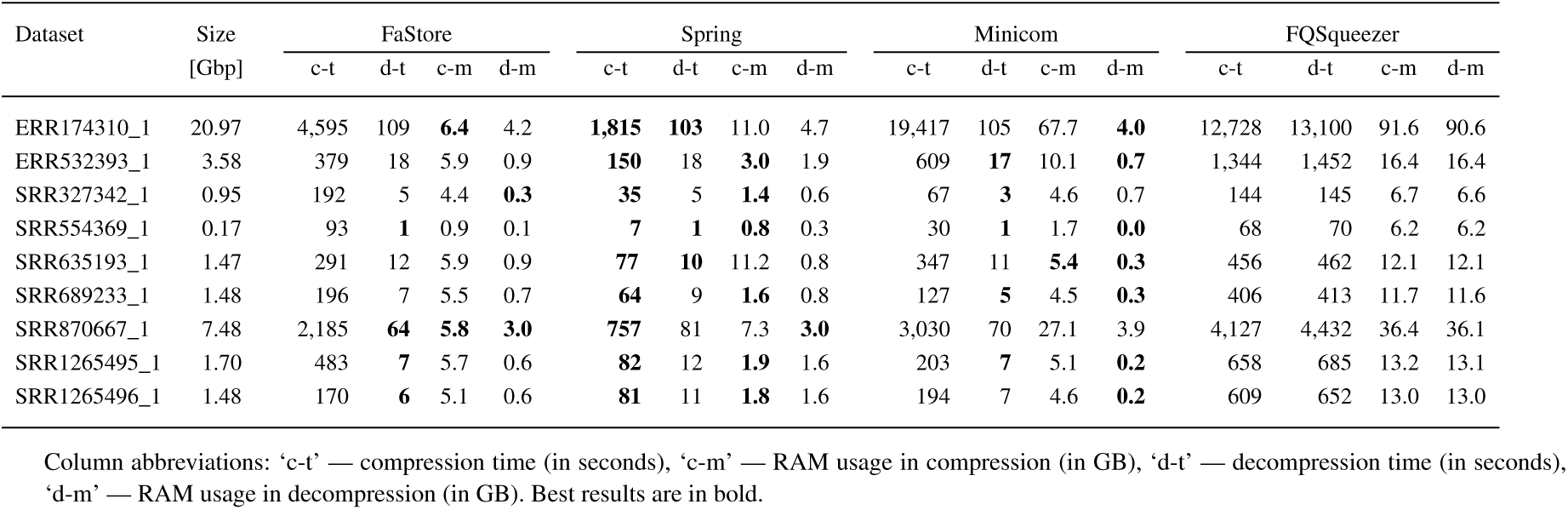
Time and memory requirements for compression of SE reads in the reordering mode

The case of memory usage is similar. The same dictionaries must be maintained by FQSqueezer in the decompression that are necessary in the compression. Moreover, to predict the successive symbols a lot of statistics must be collected. Nevertheless, without the applied correction mechanisms the occupied memory would likely be doubled.

The global dictionaries are updated only at the synchronization points, so the number of threads has some impact on the compression ratio. The results in Table 5 show that reducing the number of threads from 12 to 1 we can gain 1–2% in ratio, but the processing would be significantly slower. The last experiment in this part was to check the impact of the parameters. The results presented in Table 6 show that declaring improper genome size deteriorates the compression ratio, but the differences are not large.

**Table 5.** Multithreding scalability of FQSqueezer for SRR327342_1 dataset

| No. threads | Original ordering |  | Reordered |  |
| --- | --- | --- | --- | --- |
|  | Time [s] | Ratio [bpb] | Time [s] | Ratio [bpb] |
| 1 | 1034.76 | 0.484 | 547.72 | 0.139 |
| 2 | 619.97 | 0.485 | 378.87 | 0.140 |
| 3 | 453.10 | 0.485 | 315.44 | 0.140 |
| 4 | 356.98 | 0.486 | 263.28 | 0.141 |
| 6 | 265.40 | 0.487 | 202.04 | 0.141 |
| 8 | 221.37 | 0.487 | 168.96 | 0.142 |
| 12 | 169.56 | 0.488 | 140.31 | 0.142 |
| 16 | 143.78 | 0.489 | 131.10 | 0.143 |
| 24 | 116.33 | 0.490 | 122.69 | 0.144 |

**Table 6.**
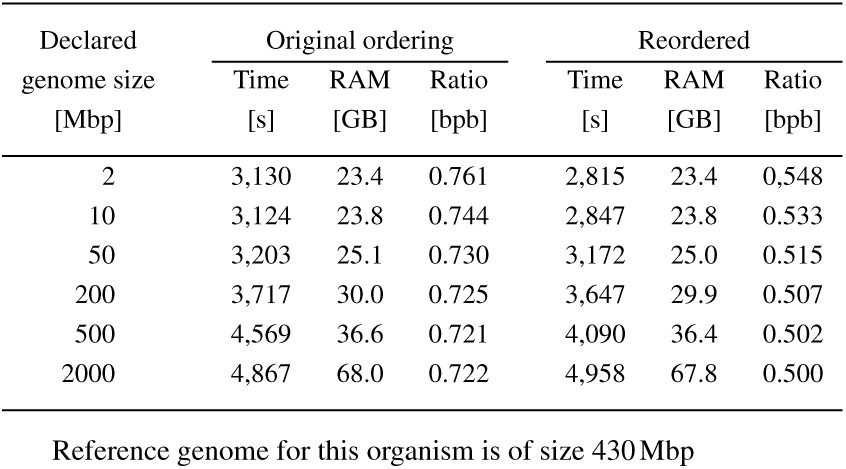
Impact of FQSqueezer declared genome size for SRR870667_1 dataset

| Declared genome size<br>[Mbp] | Original ordering |  |  | Reordered |  |  |
| --- | --- | --- | --- | --- | --- | --- |
|  | Time [s] | RAM [GB] | Ratio [bpb] | Time [s] | RAM [GB] | Ratio [bpb] |
| 2 | 3,130 | 23.4 | 0.761 | 2,815 | 23.4 | 0.548 |
| 10 | 3,124 | 23.8 | 0.744 | 2,847 | 23.8 | 0.533 |
| 50 | 3,203 | 25.1 | 0.730 | 3,172 | 25.0 | 0.515 |
| 200 | 3,717 | 30.0 | 0.725 | 3,647 | 29.9 | 0.507 |
| 500 | 4,569 | 36.6 | 0.721 | 4,090 | 36.4 | 0.502 |
| 2000 | 4,867 | 68.0 | 0.722 | 4,958 | 67.8 | 0.500 |

Reference genome for this organism is of size 430Mbp

### 3.3 Full FASTQ compressors

FQSqueezer is a FASTQ compressor, so we ran it for a few datasets to verify its performance in such situation. There were only two competitors: FaStore and Spring as Minicom was designed just for bases. The results in Table 7 are for three modes. In the *lossless* mode, all data were preserved. In the *reduced* mode, the IDs were truncated to just the instrument name and the quality values were down-sampled to 8 levels (i.e., Illumina 8-level binning). In the *bases only* mode, only the bases are stored. The table presents only the sizes of the compressed archives, but the timings and memory occupation can be found in Supplementary Worksheet.

**Table 7.** Compression of complete FASTQ files

| Dataset | Input | Original ordering |  | Reordered |  |  |
| --- | --- | --- | --- | --- | --- | --- |
|  |  | Spring | FQsqueezer | FaStore | Spring | FQsqueezer |
| Lossless |  |  |  |  |  |  |
| ERR532393_1 | 9,642 | 1,649.6 | **1,510.8** | 1,840.4 | 1,738.9 | **1,634.9** |
| SRR327342_1 | 2,813 | 435.4 | **428.2** | 504.1 | 471.5 | **468.0** |
| SRR870667_1 | 18,555 | 3,913.6 | **3,091.2** | 3,683.6 | 4,067.2 | **3,342.5** |
| ERR532393 | 19,284 | 3,200.6 | **2,903.6** | 3,602.1 | 3,299.3 | **3,032.0** |
| SRR327342 | 5,986 | 954.6 | **904.1** | — | 987.2 | **944.3** |
| SRR870667 | 32,402 | 6,060.6 | **5,093.6** | — | 6,201.6 | **5,345.0** |
| Reduced |  |  |  |  |  |  |
| ERR532393_1 | 9,642 | 917.5 | **827.8** | 978.4 | 814.4 | **723.5** |
| SRR327342_1 | 2,813 | 178.7 | **172.1** | 208.2 | 135.5 | **131.2** |
| SRR870667_1 | 18,555 | 2,583.7 | **1,824.0** | 2,151.7 | 2,444.0 | **1,620.3** |
| ERR532393 | 19,284 | 1,814.9 | **1,616.5** | 2,058.3 | 1,729.8 | **1,515.9** |
| SRR327342 | 5,986 | 413.4 | **368.9** | — | 366.6 | **328.4** |
| SRR870667 | 32,402 | 3,914.7 | **2,994.2** | — | 3,698.5 | **2,790.2** |
| Bases only |  |  |  |  |  |  |
| ERR532393_1 | 9,642 | 298.0 | **236.7** | 215.5 | 193.3 | **132.1** |
| SRR327342_1 | 2,813 | 62.1 | **59.2** | 31.6 | 18.8 | **17.0** |
| SRR870667_1 | 18,555 | 1,364.2 | **673.6** | 674.8 | 1,207.5 | **473.2** |
| ERR532393 | 19,284 | 556.6 | **416.5** | 620.6 | 451.5 | **313.5** |
| SRR327342 | 5,986 | 127.1 | **97.8** | — | 79.7 | **52.7** |
| SRR870667 | 32,402 | 1,684.5 | **871.6** | — | 1,461.0 | **667.2** |

The dataset names suffixed ‘\_1’ denote SE data. The remaining are PE files. All numbers are sizes in MBs. The best results are in bold.

The experiment confirmed the advantage of FQSqueezer in terms of compression ratio. Nevertheless, our compressor is slower and needs more memory than the competitors. It is interesting to note that in the lossless modes both Spring and FQSqueezer give smaller archives when they do not permute the reads. This is caused by the large cost of compression of IDs that are not so similar for subsequent reordered reads. In the remaining modes, the cost of ID storage is negligible, so the reordering modes win.

## 4 Conclusions

In the article, we presented a novel compression algorithm for FASTQ files. Its architecture was motivated by the PPM and DMC general-purpose compressors. Nevertheless, significant amount of work was necessary to make it possible to adapt these two approaches for genomic data. First, it was crucial to prepare specialized data structures for statistics gathering, with care of fast memory accesses (i.e., reduction of cache misses). Second, some dedicated correction of bases was implemented for better prediction and reduction of memory. Third, a special approach was necessary to efficiently find related signatures in paired reads. Fourth, the DMC-like mechanism for aggressive estimation of symbol occurrence probabilities was invented.

The experiments show advantage of FQSqueezer in term of compression ratio for majority of datasets. The differences between our tool and the state-of-the-art competitors were sometimes quite large. Nevertheless, for the largest dataset we perform slightly worse than Spring. This phenomenon deserves further investigation.

The most important drawbacks of FQSqueezer are slow processing and large memory consumption. These features are typical for PPM-based algorithms. Nevertheless, some work to reduce these drawbacks is probably possible. For example, the two most important components responsible for slow processing are looking for approximate matches and queries for incomplete *k*-mers. In the future work, it should be possible to attack at least these two problems, e.g., try to minimize the number of queries to the dictionary data structures without deterioration of the compression ratios.

## Supporting information

Supplementary Worksheet

## Supplementary material for article: FQSqueezer: *k*-mer-based compression of sequencing data

### 1 Examined programs

The following programs were used in the experimental part. The running parameters are also given.

- FaStore v. 0.8: FaStore was downloaded from https://github.com/refresh-bio/FaStore. – SE compression of bases: `./fastore_compress.sh --max --in in.fastq --threads 12 --out out.fastore` – SE decompression of bases: `./fastore_decompress.sh --in out.fastore --threads 12 --out decomp.fastq` – PE compression of bases: `./fastore_compress.sh --max --in in1.fastq --pair in2.fastq --threads 12 --out out.fastore` – PE decompression of bases: `./fastore_decompress.sh --in comp.fastore --threads 12 --out decomp.fastq` – SE compression in reduced mode: `./fastore_compress.sh --reduced --threads 12 --in in.fastq --out out.fastore` – SE compression in lossless mode: `./fastore_compress.sh --lossless --threads 12 --in in.fastq --out out.fastore` – PE compression in reduced mode: `./fastore_compress.sh --reduced --threads 12 --in in1.fastq --pair in2.fastq --out out.fastore` – PE compression in lossless mode: `./fastore_compress.sh --lossless --threads 12 --in in1.fastq --pair in2.fastq --out out.fastore`
- Spring (version from date 2018-Jul-11): Spring was downloaded from https://github.com/shubhamchandak94/Spring. – SE compression of bases, original ordering `.spring -c --no-ids --no-quality -i in.fastq -o comp.spring -t 12` – PE compression of bases, original ordering `./spring -c --no-ids --no-quality -i in1.fastq in2.fastq -o comp.spring -t 12` – SE compression of bases, reordered `./spring -c --no-ids --no-quality -i in.fastq -o comp.spring -t 12 -r` – PE compression of bases, reordered `./spring -c --no-ids --no-quality -i in1.fastq in2.fastq -o comp.spring -t 12 -r` – SE compression, lossless, original ordering `./spring -c -q lossless -t 12 -o output.spring -i in.fastq` – SE compression, lossless, reordered `./spring -c -q lossless -t 12 -o output.spring -i in.fastq -r` – PE compression, lossless, original ordering `./spring -c -q lossless -t 12 -o output.spring -i in1.fastq in2.fastq` – PE compression, lossless, reordered `./spring -c -q lossless -t 12 -o output.spring -i in1.fastq in2.fastq -r` – SE compression, reduced, original ordering `./spring -c -q ill_bin -t 12 -o output.spring -i in.fastq --no-ids` – SE compression, reduced, reordered `./spring -c -q ill_bin -t 12 -o output.spring -i in.fastq -r --no-ids` – PE compression, reduced, original ordering `./spring -c -q ill_bin -t 12 -o output.spring -i in1.fastq in2.fastq --no-ids` – PE compression, reduced, reordered `./spring -c -q ill_bin -t 12 -o output.spring -i in1.fastq in2.fastq -r --no-ids`
- Minicom (version from date 2018-Dec-27): Minicom was downloaded from https://github.com/yuansliu/minicom – SE compression of bases, reordered `./minicom -r in.fastq -t 12` – SE compression of bases, original ordering `./minicom -r in.fastq -t 12 -p` – PE compression of bases, reordered `./minicom -1 in1.fastq -2 in2.fastq -t 12`
- FQSqueezer v. 0.1: FQSqueezer was downloaded from https://github.com/refresh-bio/fqsqueezer – SE compression of bases, original ordering `./fqs-0.1 e -im n -qm n -om o -gs <genome_size> -s -t 12 -out comp.fqs in.fastq` – SE compression of bases, reordered `./fqs-0.1 e -im n -qm n -om s -gs <genome_size> -s -t 12 -out comp.fqs in.fastq` – PE compression of bases, original ordering `./fqs-0.1 e -im n -qm n -om o -gs <genome_size> -p -t 12 -out comp.fqs in1.fastq in2.fastq` – PE compression of bases, reordered `./fqs-0.1 e -im n -qm n -om s -gs <genome_size> -p -t 12 -out comp.fqs in1.fastq in2.fastq` – SE compression, lossless, original ordering `./fqs-0.1 e -gs <genome_size> -im o -qm o -om o -s -t 12 -out comp.fqs in.fastq` – SE compression, lossless, reordered `./fqs-0.1 e -gs <genome_size> -im o -qm o -om s -s -t 12 -out comp.fqs in.fastq` – PE compression, lossless, original ordering `./fqs-0.1 e -gs <genome_size> -im o -qm o -om o -p -t 12 -out comp.fqs in1.fastq in2.fastq` – PE compression, lossless, reordered `./fqs-0.1 e -gs <genome_size> -im o -qm o -om s -p -t 12 -out comp.fqs in1.fastq in2.fastq` – SE compression, reduced, original ordering `./fqs-0.1 e -gs <genome_size> -im i -qm 8 -om o -s -t 12 -out comp.fqs in.fastq` – SE compression, reduced, reordered `./fqs-0.1 e -gs <genome_size> -im i -qm 8 -om s -s -t 12 -out comp.fqs in.fastq` – PE compression, reduced, original ordering `./fqs-0.1 e -gs <genome_size> -im i -qm 8 -om o -p -t 12 -out comp.fqs in1.fastq in2.fastq` – PE compression, reduced, reordered `./fqs-0.1 e -gs <genome_size> -im i -qm 8 -om s -p -t 12 -out comp.fqs in1.fastq in2.fastq`

### 2 Datasets

**Table 1:**
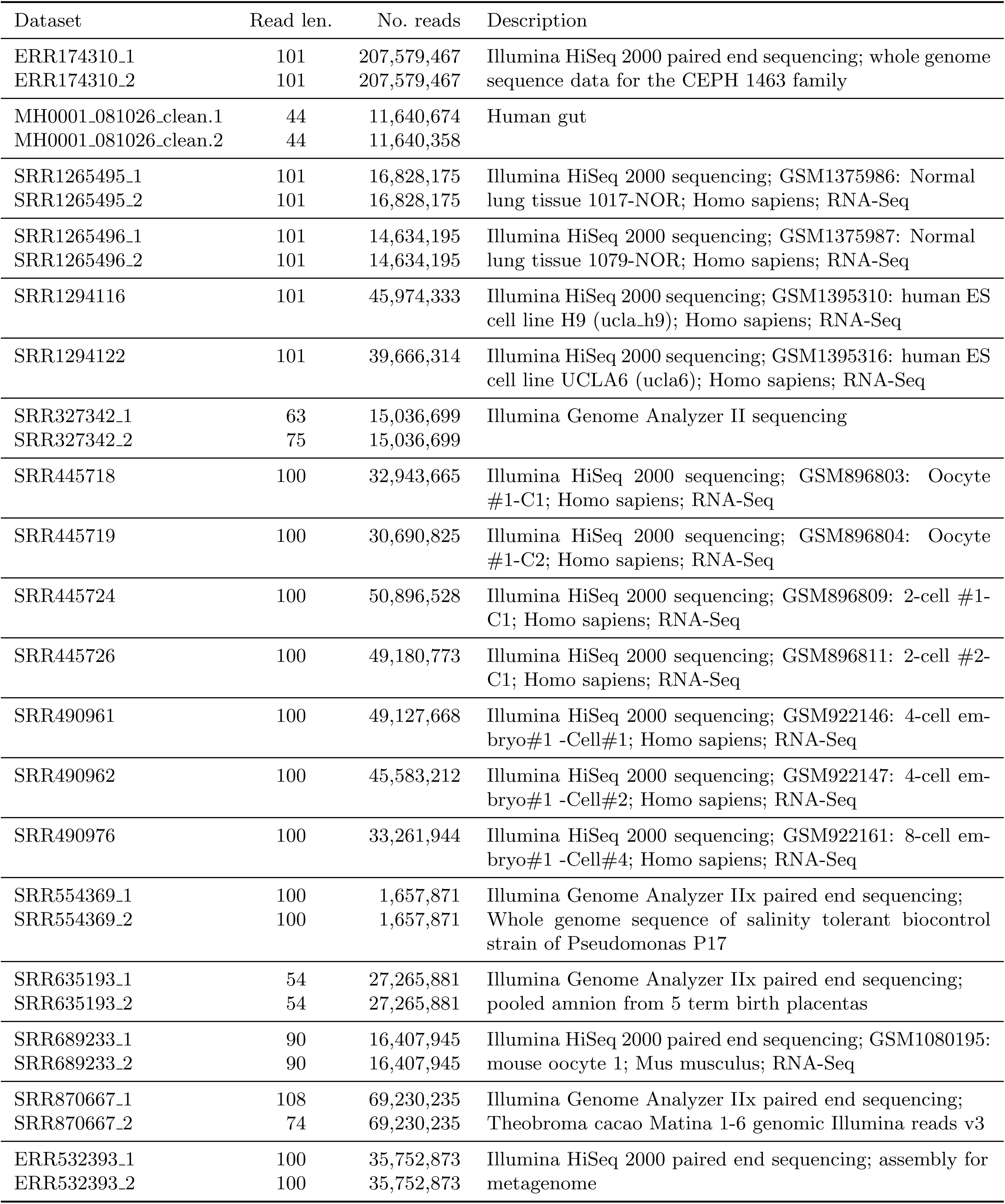
Datasets used in the experiments. The paired-end datasets were obtained from two files, i.e., ‘_1’ and ‘_2’. Descriptions are from https://www.ebi.ac.uk

### 3 Environment

The computer used in the tests was of the following configuration:

- 2 Intel Xeon E5-2670 v3 CPUs, 12 double-threaded cores per CPU, each clocked at 2.3 GHz,
- 256 GiB RAM,
- 6 HDDs of size 1 TiB each in RAID-5 configuration, `hdparm -t` reported buffered read speed 528.45 MB/s.

For compilation we used G++ v. 7.2.0. The machine was running Debian 9.3.1 x86-64 OS.

## References

Bonfield, J.K. et al. (2018) Crumble: reference free lossy compression of sequence quality values. Bioinformatics DOI: 0.1093/bioinformatics/bty608.

Bonfield, J.K. and Mahoney, M.V. (2013) Compression of FASTQ and SAM format sequencing data. PLoS One 8: e59190.

Chandak, S. et al. (2018) Compression of genomic sequencing reads via hash-based reordering: algorithm and analysis. Bioinformatics 34(4): 558–567.

Chandak, S. et al. (2018) SPRING: A next-generation compressor for FASTQ data. Bioinformatics DOI: 10.1093/bioinformatics/bty1015.

Cleary, J.G. and Witten, I.H. (1984) Data compression using adaptive coding and partial string matching. IEEE Trans. on Communications COM-32: 396–402.

Cock, P.J. et al. (2010) The Sanger FASTQ file format for sequences with quality scores, and the Solexa/Illumina FASTQ variants. Nucleic Acids Res. 38: 1767–1771.

Cormack, G.V. and Horspool, R.N.S. Data compression using dynamic Markov modelling. The Computer Journal 30: 541–550.

Cox, A.J. et al. (2012) Large-scale compression of genomic sequence databases with the Burrows–Wheeler transform. Bioinformatics 28: 1415–1419.

Deorowicz, S. et al. (2015) KMC 2: Fast and resource-frugal k-mer counting. Bioinformatics 31: 1569–1576.

Deorowicz, S. and Grabowski, S. (2011) Compression of DNA sequence reads in FASTQ format. Bioinformatics 27: 860–862.

Deorowicz, S. and Grabowski, Sz. (2013) Data compression for sequencing data, Algorithms for Molecular Biology 8: 25.

Ginart, A. et al. (2018) Optimal compressed representation of high throughput sequence data via light assembly Nature Communications 9: 566.

Grabowski, S. et al. (2015) Disk-based compression of data from genome sequencing. Bioinformatics 31: 1389–1395.

Hach, F. et al. (2012) SCALCE: boosting sequence compression algorithms using locally consistent encoding. Bioinformatics 28: 3051–3057.

Hernaez, M. et al. (2016) A cluster-based approach to compression of quality scores. In: Bilgin, A. et al. (ed.), Proc. of Data Compression Conference. IEEE Computer Society, Los Alamitos, CA, pp. 261–270.

Jones, D.C. et al. (2012) Compression of next-generation sequencing reads aided by highly efficient de novo assembly. Nucleic Acids Res., 40: e171.

Liu, Y. et al. (2018) Index suffix-prefix overlaps by (w; k)-minimizer to generate long contigs for reads compression. Bioinformatics DOI: 10.1093/bioinformatics/bty936.

Malysa, G. et al. (2015) QVZ: lossy compression of quality scores. Bioinformatics, 31, 3122–3129.

McCabe, J. (1965) On serial files with relocatable records. Operations Res. 12: 609–618.

Moffat, A. (1990) Implementing the PPM data compression scheme. IEEE Trans. on Communications, COM-38: 1917–1921.

Numanagić, I. et al. (2016) Comparison of high-throughput sequencing data compression tools. Nat. Methods 13 1005–1008.

Ochoa, I. et al. (2016) Effect of lossy compression of quality scores on variant calling. Brief. Bioinformatics 18: 183–194.

Patro, R. and Kingsford, C. (2015) Data-dependent bucketing improves reference-free compression of sequencing reads. Bioinformatics 31 2770–2777.

Roberts, M. et al. (2004) Reducing storage requirements for biological sequence comparison. Bioinformatics, 20: 3363–3369.

Roguski, L. et al. (2018) FaStore: a space-saving solution for raw sequencing data. Bioinformatics 34(16): 2748–2756.

Roguski, L. and Deorowicz, S. (2014) DSRC 2—Industry-oriented compression of FASTQ files. Bioinformatics 30: 2213–2215.

Stephens, Z.D. et al. (2015) Big Data: astronomical or genomical. PLoS Biol. 13: e1002195.

